# A genetically validated approach to detect inorganic polyphosphates in plants

**DOI:** 10.1101/630129

**Authors:** Jinsheng Zhu, Sylvain Loubéry, Larissa Broger, Laura Lorenzo-Orts, Anne Utz-Pugin, Chang Young-Tae, Michael Hothorn

## Abstract

Inorganic polyphosphates (polyPs) are linear polymers of orthophosphate units linked by phosphoanhydride bonds. PolyPs represent important stores of phosphate and energy, and are abundantly found in many pro- and eukaryotic organisms. In plants, the existence of polyPs has been established using microscopy and biochemical extraction methods that are now known to produce artifacts. Here we use a polyP-specific dye and a polyP binding domain to detect polyPs in plant and algal cells. To develop the staining protocol, we induced polyP granules in *Nicotiana benthamiana* and Arabiopsis cells by heterologous expression of *E. coli* polyphosphate kinase 1 (PPK1). Over-expression of PPK1 but not of a catalytically impaired version of the enzyme lead to severe growth phenotypes, suggesting that ATP-dependent synthesis and accumulation of polyPs in the plant cytosol is toxic. We next crossed stable PPK1 expressing Arabidopsis lines with plants expressing the polyP-binding domain of *E. coli* exopolyphosphatase (PPX1c), which co-localized with PPK1-generated polyP granules. These granules were stained by the polyP-specific dye JC-D7 and appeared as electron dense structures in transmission electron microscopy (TEM) sections. Using the polyP staining protocol derived from these experiments, we screened for polyP stores in different organs and tissues of both mono- and dicotyledonous plants. While we could not detect polyP granules in higher plants, we could visualize the polyP-rich acidocalicisomes in the green algae *Chlamydomonas reinhardtii.* Together, our experiments suggest that higher plants may not contain large polyPs stores.

**Significance Statement:** A chemical dye and an inorganic polyphosphate binding domain are shown to specifically label inorganic polyphosphate granules in transgenic Arabidopsis lines and Chlamydomonas acidocalcisomes. Using these tools, we show that in contrast to many prokaryotic and eukaryotic organisms, higher plants do not seem to contain large inorganic polyphosphate stores.

## Introduction

PolyPs are linear inorganic phosphate (Pi) polymers, with sizes ranging from tripolyphosphate (3 Pi units) to long-chain polyPs (~1,000 Pi units) (Kornberg, 1956; Kornberg *et al.*, 1957; Kulaev and Vagabov, 1983; Clark and Wood, 1987). In bacteria, polyPs accumulate in volutin granules (Babes, 1895; Widra, 1959; Tumlirsch and Jendrossek, 2017) or in membrane-surrounded acidocalcisomes (Seufferheld *et al.*, 2003). Bacterial polyPs are synthesized by two different classes of polyphosphate kinases PPK1 (Kornberg, 1957; Ahn and Kornberg, 1990) and PPK2 (Zhang *et al.*, 2002) from ATP or GTP, respectively. In yeast and other fungi, large amounts of long-chain polyPs (Macfarlane, 1936) generated by the vacuolar transporter chaperone (VTC) complex (Hothorn *et al.*, 2009) accumulate in the vacuole (Urech *et al.*, 1978) and in the cell wall (Werner *et al.*, 2007b). The eukaryotic VTC complex is conserved among other unicellular eukaryotes such as protozoa (Fang *et al.*, 2007) and green algae (Aksoy *et al.*, 2014), organisms that store polyPs in specialized vacuoles termed acidocalcisomes (Docampo and Huang, 2016) and also in their cell walls (Werner *et al.*, 2007a). The eukaryotic amoeba *Dictyostelium discoideum* contains polyP accumulating acidocalcisomes but lacks the VTC complex. Instead *Dictyostelium* features two polyphosphate kinases that are similar to bacterial PPK1 (Zhang *et al.*, 2005) and to an actin-related protein (Gómez-García and Kornberg, 2004), respectively. Neither PPK1, PPK2 nor the VTC complex appear to be present in higher eukaryotes (Hooley *et al*., 2008), but polyPs of various chain-length have been identified in mitochondria and lysosomes of different animal and human cells and tissues (Pisoni and Lindley, 1992; Kumble and Kornberg, 1995). PolyPs also accumulate in acidocalcisomes in human platelet and mast cells (Ruiz *et al.*, 2004; Moreno-Sanchez *et al.*, 2012).

Invasive and non-invasive polyP detection techniques have been developed and applied to detect specific polyP stores in bacteria, fungi and algae, all of which accumulate polyP to high levels. Initially, polyP bodies were detected in microorganisms as metachromatic granules in light microscopy experiments, and stained with unspecific dyes such as toluidine blue (Meyer, 1904). Using this method, polyP granules were reported from lower plants, including algae and different mosses (Keck and Stich, 1957). In TEM sections, electron dense granules have been interpreted as polyP bodies or acidocalcisomes, for example in the seeds of different palm species and in rice anthers (DeMason and Stillman, 1986; Mamun *et al.*, 2005).

PolyPs can also be biochemically purified and initially the acid-labile polymer was converted to Pi using strong acids. This rather unspecific and insensitive method was used early on to characterize polyP stores in spinach leaves and in parasitic plants (Miyachi, 1961; Tewari and Singh, 1964). The identification of highly specific bacterial and fungal polyP metabolizing enzymes, such as *E. coli* PPK1 (which can also catalyze the reverse reaction, the phosphorylation of ADP to ATP using polyP as a phosphate donor) or the *E. coli* (Akiyama *et al.*, 1993) and *S. cerevisiae* (Wurst and Kornberg, 1994) exopolyphosphatase PPX1, has enabled the enzymatic identification and quantification of polyPs extracted from cells and tissues (Ault-Riché and Kornberg, 1999; Bru *et al.*, 2016). This method has, to the best of our knowledge, not been applied to study polyP stores in higher plants.

The isolated polyP-binding domain of *E. coli* PPX1 has been used to immunolocalize polyPs in fungal, algae and human cells (Saito *et al.*, 2005; Werner *et al.*, 2007b; Werner *et al.*, 2007a; Jimenez-Nuñez *et al.*, 2012). PolyPs of different chain length can also be visualized on UREA-PAGE gels, either using radioactive ^32^P, toluidine blue or 4’,6-diamidino-2-phenylindol (DAPI) (Ogawa *et al.*, 2000; Smith and Morrissey, 2007). *In vivo* ^31^P nuclear magnetic resonance spectroscopy has been used to define phosphate-containing metabolites in cells and tissues, including polyP in bacteria and lower eukaryotes (Moreno *et al.*, 2000). Raman spectroscopy has been used to detect Pi-rich inositol polyphosphates, but not inorganic polyphosphates in wheat grains (Kolozsvari *et al.*, 2015). *In situ* staining methods for polyP have been developed using DAPI (Kulakova *et al*., 2011). Its unique emission spectrum allows to distinguish polyP from nucleic acids (also interacting with DAPI), but not from inositol polyphosphates also abundant in plants (Kolozsvari *et al.*, 2014). Two specific dyes for polyP, JC-D7 and JC-D8 have been developed and employed to localize polyP stores in mammalian cells.

We have recently uncovered a molecular connection between the Pi starvation response controlled by inositol pyrophosphate (PP-InsPs) signaling molecules and their SPX receptors, and polyP metabolism in yeast (Wild *et al.*, 2016). PP-InsPs and SPX domains also control the Pi starvation response in Arabidopsis (Zhu *et al.*, 2018), and we have characterized the short-chain polyphosphatase TRIPHOSPHATE TUNNEL METALLOENZYME 3 (AtTTM3) (Martinez *et al.*, 2015; Lorenzo-Orts, Witthoeft, *et al.*, 2019) and a specific polyP-binding domain (Lorenzo-Orts, Hohmann, *et al.*, 2019) in plants. We thus speculate that polyPs may exist in plants and that they could form a relevant Pi store. Here we make use of both the JC-D7 dye (Angelova *et al.*, 2014) and the PPX1 polyP-binding domain to probe for the presence of polyP stores in different plant cells and tissues.

## Results

To test if JC-D7 (Angelova *et al.*, 2014) can label polyP in intact plant cells we sought to introduce defined polyP stores, allowing for a genetically validated detection of polyP. To this end, we generated transgenic Arabidopsis lines expressing the bacterial polyphosphate kinase PPK1 from *E. coli,* which polymerizes long-chain polyPs from ATP (Kornberg, 1957; Ahn and Kornberg, 1990), under the control of the Ubi10 promoter and carrying a C-terminal mCitrine (mCit) tag (see Methods). EcPPK1 has been previously characterized as a histidine kinase, and mutation of His435 to alanine impaired polyP accumlation in *E. coli* (Kumble *et al.*, 1996). The crystal structure of EcPPK1 in complex with a non-hydrolyzable substrate analog revealed that His435 and His592 are in direct contact with the β- and γ-phosphates of the ATP substrate (Figure 1a) (Zhu *et al.*, 2005). Based on this analysis, we generated a control line expressing a catalytically impaired PPK1 (mPPK1), in which His435 and His592 are replaced by alanine. PPK1-mCit T3 lines displayed a strong growth phenotype with small purple leaves, which correlates with PPK1-mCit protein levels (Figure 1b,c). The mPPK1-mCit control lines are similar to wild-type.

**Figure 1.**
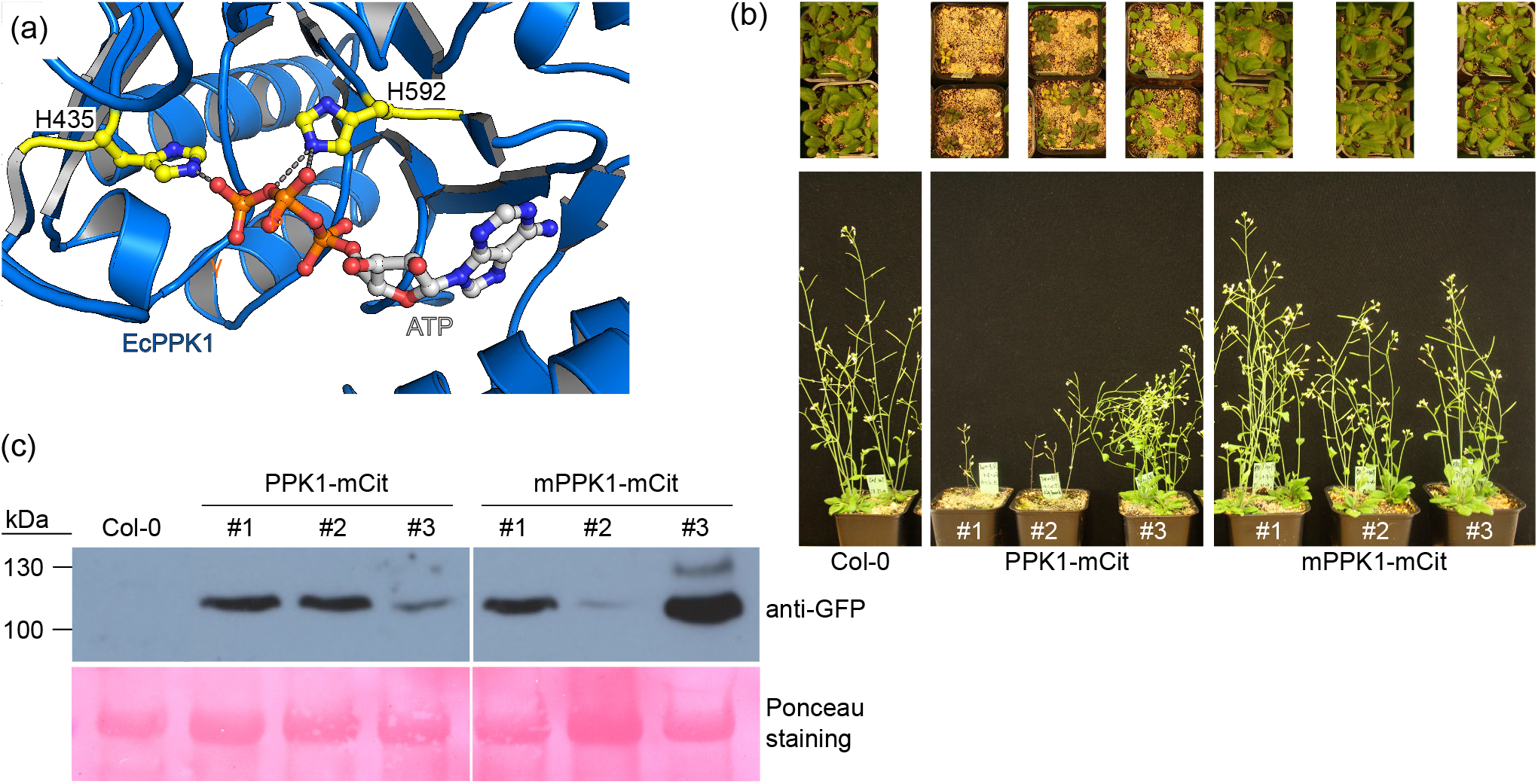
Heterologous expression of *E. coli* PPK1 results in dose-dependent stunted growth in Arabidosis. (a) Structural view of the PPK active site. PPK1 is show in blue (PDB-ID 1XDO, ribbon diagram) with the catalytic His435 and His592 (in bonds representation, in yellow) contacting the γ and β phosphates of the ATP substrate (in bonds representation). (b) Growth phenotype of PPK1 and mPPK1 expressing plants. Shown are rosettes of 4 week old plants (upper panel) and 5 week old flowering plants (lower panel). Three independent lines are shown for each PPK1-mCit or mPPK1-mCit. (c) Western blot of the lines shown in (b, lower panel), using an anti-GFP antibody.

mPPK1-mCit control plants show a uniform expression of the fusion protein in root tips (Figure 2). In contrast, expression of the catalytically active PPK1-mCit is patchy, with some cells accumulating the fusion protein to high levels and others showing small punctate structures. These punctate structures appear as dense granules in the corresponding bright field images (Figure 2).

**Figure 2.**
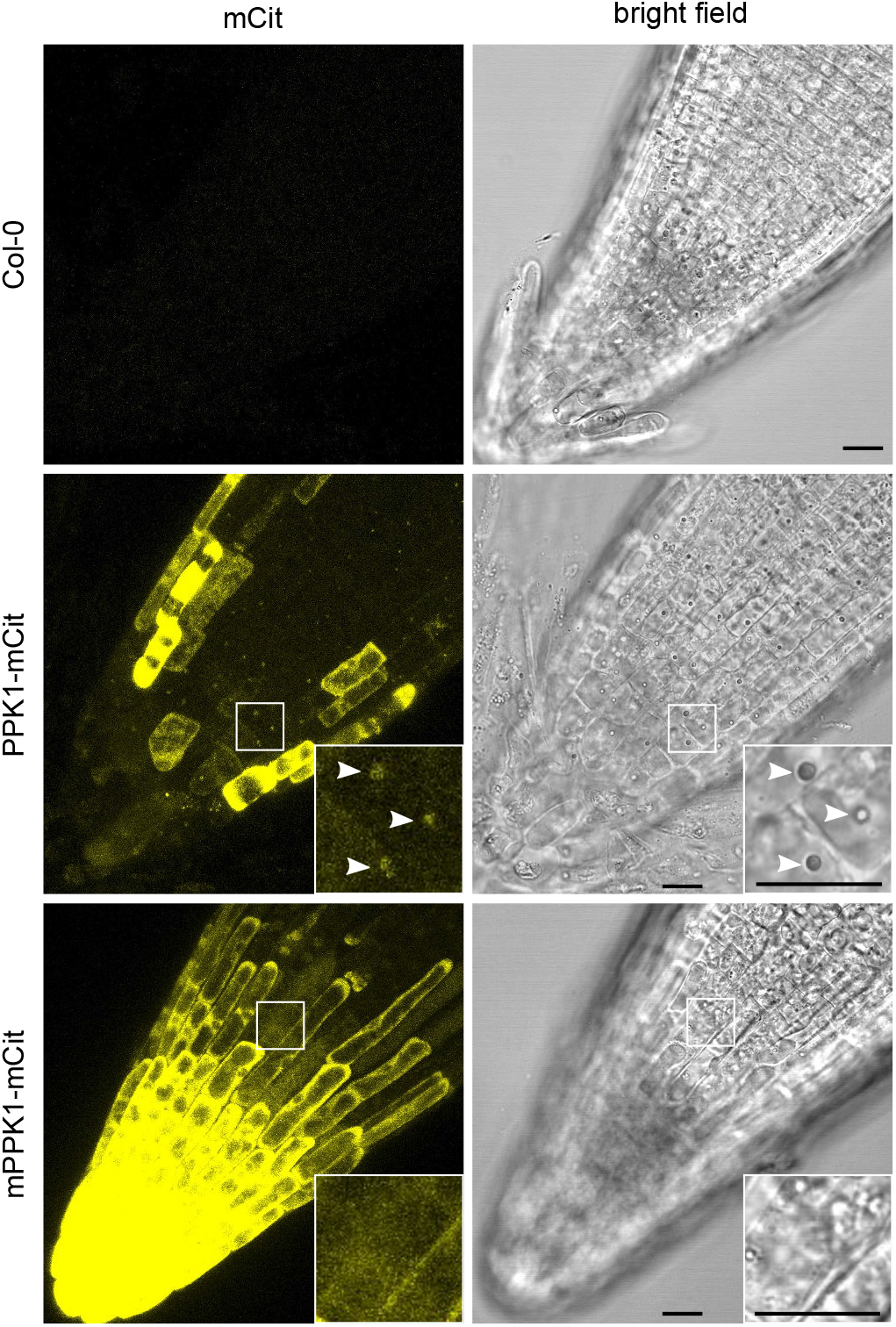
PPK1-mCit but not mPPK1-mCit expression results in the formation of punctate structures surrounded by the enzyme. Confocal imaging of root tips of 7 DAG seedlings grown on ^1/2^MS plates. Col-0, PPK1-mCit and mPPK1-mCit expressed seedlings were imaged with similar confocal settings. Enlarged squares highlight the presence of dense punctate structures (highlighted by arrow heads) in PPK1 but not mPPK1 expressing lines. Scale bar = 20 μm.

We next tested if ectopic expression of PPK1 introduces polyP stores in plant cells and if those stores can be detected using the JC-D7 dye. We transiently expressed the PPK1-mCit and mPPK1-mCit constructs in *Nicotiana benthamiana* leaves, where we again found mPPK1-mCit to be evenly distributed in the cytosol, while PPK1-mCit localized in punctate structures (Figure 3a). We stained epidermal cells with JC-D7 (see Methods), and found the dye to co-localize with the punctate structures in PPK1-mCit expressing cells, but not in the mPPK1-mCit control (Figure 3a). We hypothesized that PPK1-mCit localized to polyP granules generated by the bacterial enzyme, which are labeled by JC-D7. To test this hypothesis, we used the polyP binding domain of the *E. coli* exopolyphophatase PPX1 (Akiyama *et al.*, 1993), which has been previously employed to specifically stain polyP stores in fungal, algal and human cells (Saito *et al.*, 2005; Werner *et al.*, 2007a; Werner *et al.*, 2007b; Jimenez-Nuñez *et al.*, 2012). To this end, we generated a PPXc – mCherry (mChe) fusion protein (residues 311-508; Figure 3b). We found PPXc-mChe to localize to the cytoplasm and nucleus when transiently expressed in *Nicotiana benthamiana*, and in stable transgenic Arabidopsis lines (Figure 3c). Next, we crossed PPX1c-mChe with our PPK1-mCit and mPPK1-mCit lines, which revealed PPXc-mChe to co-localize with PPK1-mCit in JC-D7 stained granules in hypocotyl cells (Figure 3d). This was not observed in the mPPK1-mCit control lines (Figure 3d). Co-localization of PPK1-mCit, PPXc-mChe and the JC-D7 dye could also be observed in fixed leaf epidermal and hypocotyl cells in Arabidopsis (Figure S1). TEM analysis of hypocotyl sections revealed the presence of electron-dense granules in PPK1-mCit but not in mPPK1-mCit expressing lines (Figure 4). Together, these experiments suggest that PPK1 introduces polyP granules in Arabidopsis, which can be detected by both the JC-D7 dye and the PPX1c polyP-binding domain.

**Figure 3.**
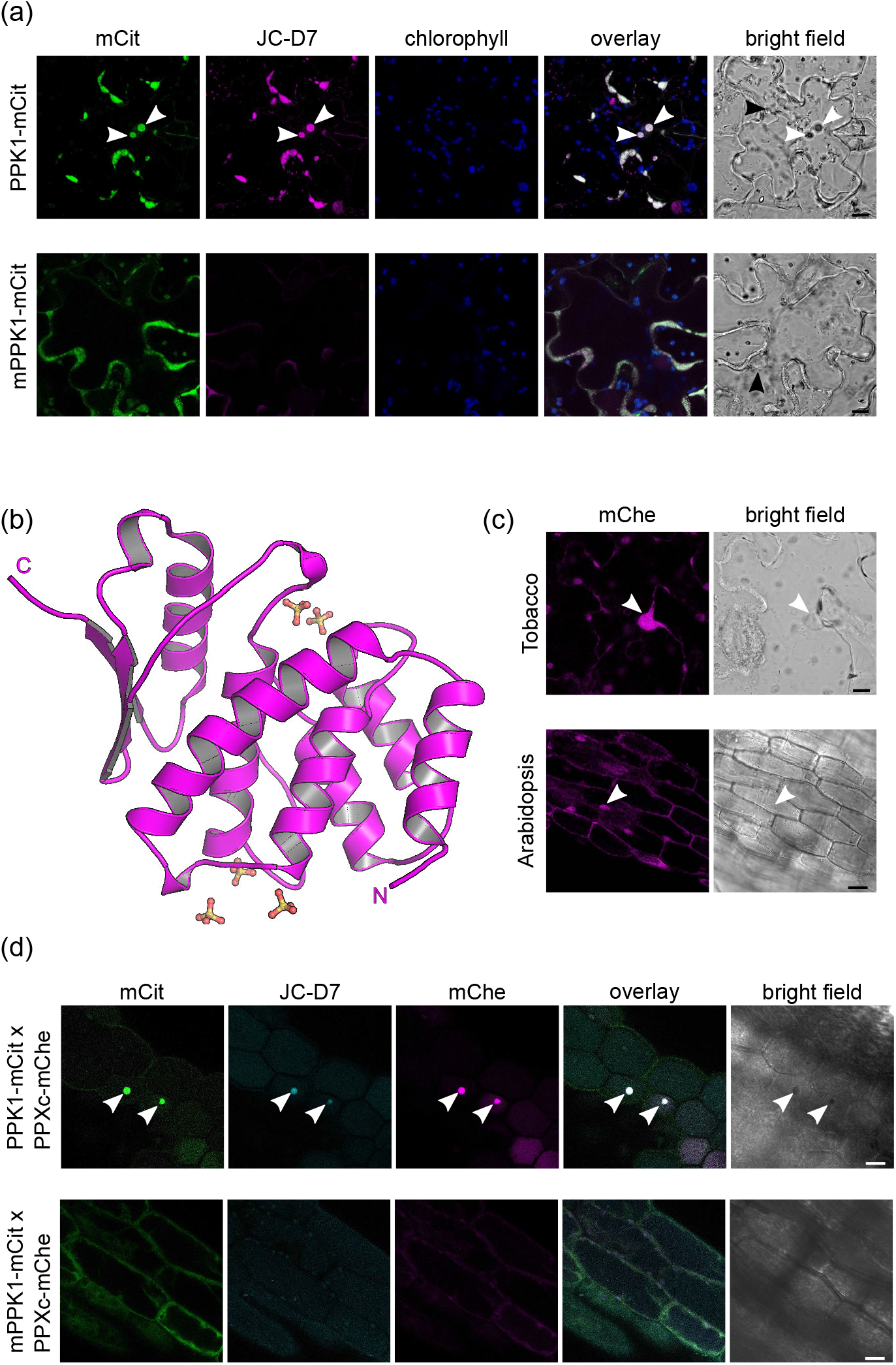
JC-D7 and PPXc mark polyP granules in PPK1-mCit but not mPPK1-mCit expressing plants. (a) Transiently expressed PPK1-mCit (green) co-localizes with JC-D7 (magenta) stained polyP granules outside chloroplasts (blue) in tobacco epidermal cells. White arrow heads highlight the granule structures, and black arrow heads indicate putative nuclear structures. (b) Overview of the PPX polyP binding domain PPXc (PDB-ID 1U6Z, residues 311-508, ribbon diagram, in magenta) bound to sulfate ions which mimic the position of the polyP substrate (Alvarado *et al.*, 2006) (in bonds representation). (c) Sub-cellular localization of mChe tagged PPXc (magenta) expressed in isolation in *Nicotiana benthamiana* epidermal cells and Arabidopsis hypocotyl cells. Arrow heads indicate putative nuclear structures. (d) Arabidopsis lines that stably express PPXc-mChe were crossed with lines that stably express PPK1-mCit and mPPK1-mCit. Active PPK1-mCit (green, upper panel) but not its catalytic impaired mutant version (green, lower panel) co-localizes with PPXc-mChe (magenta) on JC-D7 (cyan) stained polyP granules. Scale bars = 10 μm.

**Figure 4.**
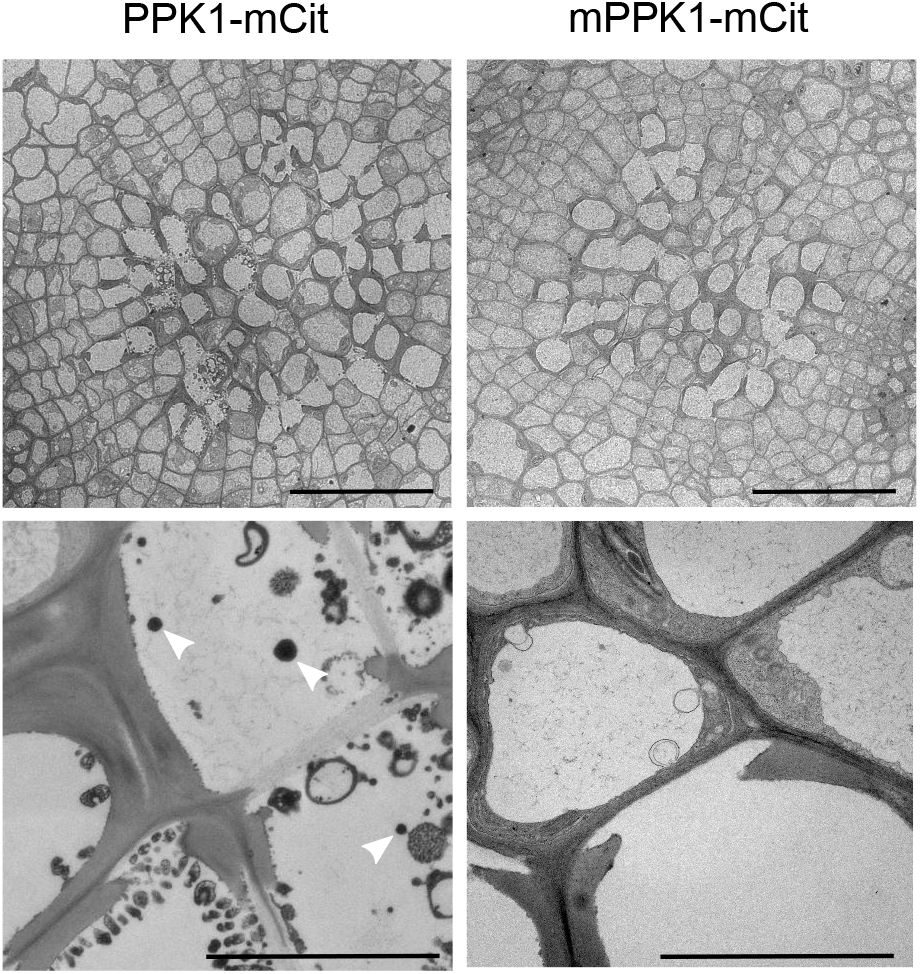
Electron dense granules are present in TEM sections of PPK1-mCit but not mPPK1-mCit expressing plants. Hypocotyl sections (upper panel, scale bar = 50 μm) together with a zoomed in view on xylem cells (lower panel, scale bar = 5 μm) of PPK1-mCit and mPPK1-mCit expressing Arabidopsis lines. Arrow heads highlight electron dense granules.

While we could readily detect PPK1-generated polyP granules in different cells and tissues from Arabidopsis and tobacco, we could not reproducibly detect JC-D7 stained structures in Col-0 wild-type (Figures S2, S3). PolyP also did not accumulate in a previously characterized loss-of-function mutant of AtTTM3 *(ttm3-1)* (Moeder *et al.*, 2013; Lorenzo-Orts, Witthoeft, *et al.*, 2019). (Figures S2, S3). We next stained different mono-and dicots using the JC-D7 dye (examples are shown in Figure S4) but again could not detect a polyP-specific signal. In contrast, JC-D7 stained the previously characterized polyP containing acidocalcisomes of *Chlamydomonas reinhardtii,* which appeared enriched in a sulfur depleted growth condition (Gal *et al.*, 2018; Aksoy *et al.*, 2014) (Figure 5).

**Figure 5.**
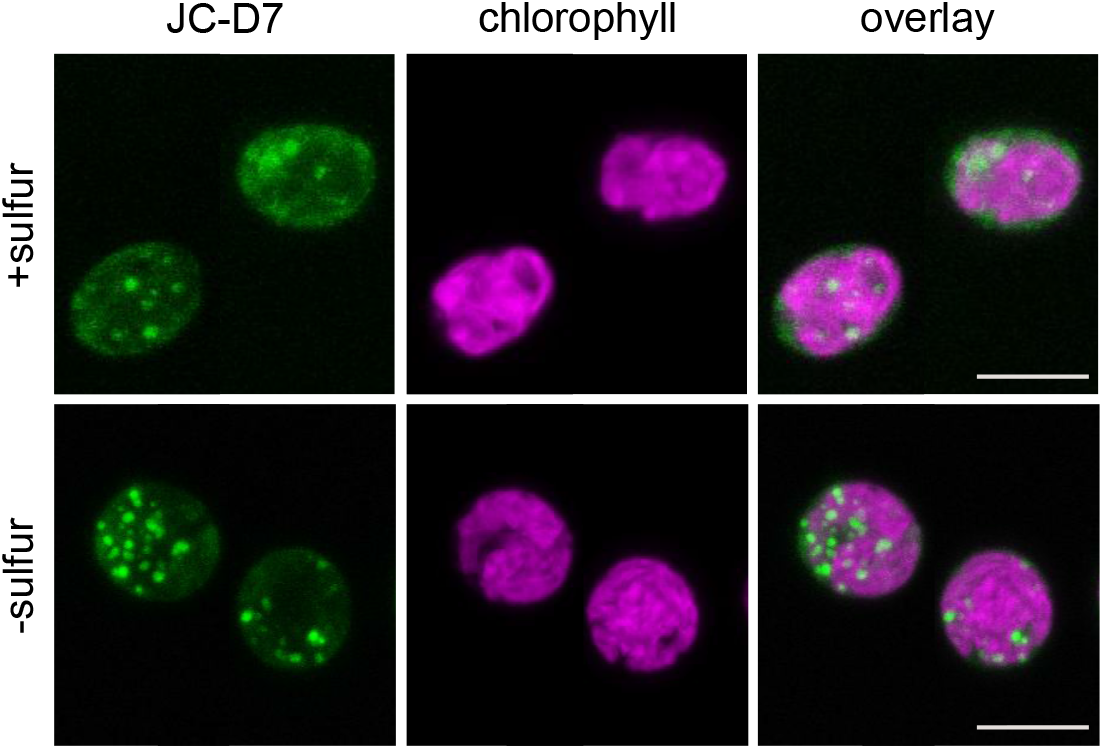
JC-D7 can specifically stain polyP granules in *Chlamydomonas reinhardtii.* JC-D7 (green) stained polyP granules in Chlamydomonas cells grown in tris-acetate phosphate medium. The-sulfur medium was prepared by substituting sulfates by their corresponding chloride salts. Chlorophyll auto-fluorescence is shown in magenta. Scale bars = 10 μm.

## Discussion

PolyPs are evolutionary ancient energy polymers and have been identified in a variety of pro- and eukaryotic organisms (Rao *et al.*, 2009). Based on earlier reports (Keck and Stich, 1957; Miyachi, 1961; Tewari and Singh, 1964; DeMason and Stillman, 1986), polyPs have been assumed to exist in higher plants (Clark and Wood, 1987; Kornberg *et al.*, 1999; Kulaev, 2005). However, just like in animals the synthesis, storage and breakdown of polyP has been poorly characterized and no physiological functions have been assigned to polyP in plants. In the green algae *Chlamydomonas reinhardtii* polyP stores have been identified in acidocalcisomes (Ruiz *et al.*, 2001; Gal *et al.*, 2018) and in the cell wall compartment (Werner *et al.*, 2007a), as it has been seen in other unicellular eukaryotes such as yeast or trypanosomes (Docampo and Huang, 2016). PolyP in Chlamydomonas is synthesized by the vacuolar transporter chaperone VTC complex (Aksoy *et al.*, 2014), whose catalytic subunit Vtc4 generates polyP from ATP (Hothorn *et al.*, 2009), and translocates the growing polymer to the vacuole/acidocalcisome using a membrane pore (Gerasimaitė *et al.*, 2014). While the VTC complex is apparently absent in higher plants, proteins sharing structural homology with the Vtc4 catalytic domain exist in Arabidopsis (Moeder *et al.*, 2013; Martinez *et al.*, 2015; Ung *et al.*, 2017). We have previously characterized AtTTM3 as a specific short-chain inorganic polyphosphatase (Martinez *et al*., 2015), which is transcribed in a bicistronic transcript also encoding the cell cycle regulator AtCDC26 (Lorenzo-Orts, Witthoeft, *et al.*, 2019).

Since the polyP-synthesizing enzymes in higher eukaryotes remain to be identified, we set out to develop a genetically validated method that would allow for the specific detection of polyP stores in living cells. DAPI has been used to visualize polyP stores in various pro- and eukaryotic cells using a unique DAPI-polyP emission spectrum (Aschar-Sobbi *et al.*, 2008; Puchkov, 2010; Kulakova *et al.*, 2011; Gomes *et al.*, 2013; Martin and Van Mooy, 2013). However, DAPI has recently been shown to also interact with inositol phosphates (Kolozsvari *et al.*, 2014) and many of the staining procedures lacked suitable genetic controls that would confirm the identity of the stained structures as *bona fide* polyP stores. We thus chose to heterologously express a bacterial polyP kinase PPK1 to induce specific polyP granules in *Nicotiana benthamiana* and Arabidopsis cells (Figures 1,3). We found that expression of PPK1 but not of a catalytically inactive version of the enzyme led to stunted growth and anthocyanin accumulation in stable transgenic lines. This is possibly due to the accumulation of polyPs in the cytosol, which has been previously shown to have severe toxic effects in yeast (Gerasimaitė *et al.*, 2014). As polyPs are very potent chelators of divalent cations, the presence of polyPs in the cytosol may inhibit the activity of many enzymatic processes and disrupt cellular structures. In line with this, we observe patchy expression of a PPK1-mCit but not the mPPK1-mCit fusion protein in the root tip of Arabidopsis (Figure 2).

The PPK1-generated polyP granules in Arabidopsis co-localize with the PPK1-mCit fusion protein itself and with the polyP-specific binding domain PPXc from *E. coli* (Figure 3). Importantly, these structures were specifically stained with the JC-D7 dye, previously shown to detect polyP granules in human cells (Angelova *et al.*, 2014). The same structures appear as electron dense granules in TEM analysis (Figure 4). Together, our experiments suggest that JC-D7 is a specific dye for polyP detection in plants. In line with this, JC-D7 stained the known, VTC-generated polyP stores in Chlamydomonas (Ruiz *et al*., 2001; Aksoy *et al*., 2014). However, we failed to reproducibly detect polyP granules in wild-type Arabidopsis, tobacco, maize, *Phaesolus vulgaris* or *Ricinus communis* plants (Figures. 3, S2–4), the latter of which contains the polyP-specific binding protein RcCHAD (Lorenzo-Orts, Hohmann, *et al.*, 2019). Also, we could not detect polyP accumulation in *ttm3-1* mutant, which we hypothesized to have reduced polyP phosphatase activity (Figures S2,S3) (Martinez *et al.*, 2015; Lorenzo-Orts, Witthoeft, *et al.*, 2019). These findings suggest that either plants do not contain polyP stores, that they are present in concentrations too low to be detected by JC-D7, or that they do not accumulate in the tissues and/or growth conditions analyzed. In line with this, inositol polyphosphates including phytic acid (inositol hexakisphosphate) but not inorganic polyPs were detected by Raman spectroscopy in wheat grains (Kolozsvari *et al.*, 2015). However, our results are in stark contrast with earlier reports relying on biochemical extraction and hydrolysis methods (Miyachi, 1961; Tewari and Singh, 1964; DeMason and Stillman, 1986), which inadvertently may have detected inositol polyphosphates and/or other organic phosphates.

Taken together, we could demonstrate that indeed JC-D7 can specifically stain dense polyP granules in living and fixed plant and algal cells, but we failed to detect significant polyP stores in wild-type mono- or dicotlyledonous plants. Of course, we cannot rule out the possibility that higher plants only accumulate polyP in specific organs or cells, only in certain developmental stages, or in response to certain environmental stimuli. The lack of significant polyP stores however raises the question, why conserved, broadly expressed polyP phosphatases exist in plants (Moeder *et al.*, 2013; Martinez *et al.*, 2015; Lorenzo-Orts, Witthoeft, *et al.*, 2019) and why some species adopted polyP-specific binding proteins (Lorenzo-Orts, Hohmann, *et al.*, 2019). At this point, despite our efforts the statement by Harold (1966) remains true: “The case for the occurrence of polyP in higher plants and animals would be materially strengthened by the demonstration of appropriate biosynthetic enzymes” (Harold, 1966).

## Experimental procedures

### Plant material and growth conditions

Plants were grown at 21°C with 50 *%* humidity and a 16 h light : 8 h dark cycle. For confocal imaging, Arabidopsis seedlings were generally grown on vertical plates with half-strength Murashige and Skoog (^1/2^MS, Duchefa) media containing 1% (w/v) sucrose, 0.5 g/L MES buffer and 0.8 % (w/v) agar (Duchefa), pH 5.7 for 7 to 8 days. Seeds were sequentially surface-sterilized using 70 % (v/v) ethanol, a 5 % hypochlorite solution (Javel, 13 – 14 %), and rinsed 4 times with sterilized water.

The Arabidopsis *ttm3-1* (SALK_133625) mutant was obtained from the European Arabidopsis Stock Center (http://arabidopsis.info/) (Lorenzo-Orts, Witthoeft, *et al.*, 2019). *Chlamydomonas reinhardtii* strain WT 2C (mt^+^) cells were grown in tris-acetate phosphate medium (Kropat *et al.*, 2011), in white light originating from fluorescent tubes at 60 μM photons m-2 s-1 and at 25°C. The-sulfur medium was prepared by substituting sulfates by their corresponding chloride salts.

### Generation of Arabidopsis transgenic lines

The coding sequences of *Escherichia coli* PPK1 (UniProt ID C3T032) and the mPPK1 catalytically inactive mutant (H435A; H592A) were assembled together with a Ubi10 promoter and a C-terminal mCit tag and introduced into destination vector pH7m34GW2 and pB7m34GW2 respectively as described (Lorenzo-Orts, Hohmann, *et al.*, 2019). The coding sequence of the C-terminal polyP binding domain of *Escherichia coli* exopolyphosphatase (PPXc, UniPro ID P0AFL6) was cloned into a pDONR221 vector using forward primer GGGGACAAGTTTGTACAAAAAAGCAGGCTTAATGCATCAGGATGTGCGTAGTCGC and reversed primer GGGGACCACTTTGTACAAGAAAGCTGGGTAAGTACTTTCTTCTTCAATTTTC. The PPXc domain was fused to a C-terminal mChe tag and expression was driven by a Ubi10 promoter, assembled in destination vector pH7m34GW2. All of the constructs were transformed into *A. tumefaciens* strain pGV2260 by heat shock. All of transgenic lines were generated in Col-0 background using the floral dip method (Clough and Bent, 1998). T3 generation transformants were selected in ^1/2^MS medium (Duchefa), supplement with 50 μg/mL BASTA or 20 μg/mL Hygromycin.

### Confocal imaging

All light microscopy imaging was performed on a Zeiss LSM780 confocal microscope using a 40 × / 1.3 water objective. Various confocal settings were set to record the emission of mCitrine (excitation 514 nm; emission 517-550 nm), mCherry (excitation 594 nm; emission 606-632nm), and JC-D7 (excitation 405 nm; emission 480-510 nm) using a GaAsP detector, and chlorophyll (excitation 594 nm; emission 653-698 nm) was imaged with a PMT detector. The sequential scanning mode was used for imaging multiple-color images. JC-D7 (Angelova *et al.*, 2014) (Glixx Laboratories Inc.) was applied at a concentration of 100 μM for 1.5 h at room temperature in HEPES buffered salt solution (HBSS) composed of 10 mM HEPES/NaOH (pH 7.4), 156 mM NaCl, 3 mM KCl, 2 mM MgSO_4_, 1.25 mM KH_2_PO_4_, 2 mM CaCl_2_, 10 mM glucose, as described (Angelova *et al.*, 2014). 50 mM stock solution were prepared in 100% (v/v) DMSO. Live cell imaging was performed in root, hypocotyl, and leaf epidermis cells. Plant tissues or *C. reinhardtii* strain WT 2C (mt^+^) were incubated together with JC-D7 in HBSS buffer. To image fixed plant tissues, samples were fixed in vacuum in PBS buffer containing 4 *%* (v/v) para-formaldehyde, 0.1 *%* (v/v) Triton X-100 and 1 mM DTT (in the case of roots), then sequentially washed two times in PBS and two times in HBSS buffer.

To co-localize PPK1-induced polyP granules with PPXc-mChe and JC-D7, Arabidopsis lines harboring PPK1-mCit or PPK1m-mCit were crossed with plants expressing PPXc-mChe. F1 generation seeds were grown on ^1/2^MS medium for 8-9 d and then stained by JC-D7 as described above. To co-localize transiently expressed PPK1-mCit and PPK1m-mCit with JC-D7, *A. tumefaciens* cell cultures were infiltrated into *Nicotiana benthamiana* leaves as described (Lorenzo-Orts, Hohmann, *et al.*, 2019). Epidermal cells were imaged by confocal microscopy after 3 d incubation at room temperature.

### Transmission electron microscope (TEM) imaging

Samples for TEM were fixed overnight at 4 °C in 2.5% (v/v) glutaraldehyde, 0.01% (v/v) Tween-20 in 100 mM sodium cacodylate pH 7.0, after vacuum infiltration. A first post-fixation was done in 1.5 % (w/v) osmium tetroxide for 2 h at 4 °C, and a second post-fixation in 1 % (w/v) uranyl acetate for 1 h at 4 °C. Seedlings were then embedded in 1.5 % (w/v) agarose, dehydrated in a graded ethanol series and finally embedded in Epon 812. 85 nm ultra-thin sections were made using a UCT microtome (Leica) and deposited on Formvar films on copper grids; they were stained for 25 min with 2 % (w/v) uranyl acetate, then for 20 min with Reynolds lead citrate, and finally observed with a Tecnai G2 Sphera (FEI) at 120kV equipped with a high-resolution digital camera.

### Plant protein extraction for western blot

Around 250 mg plant tissue was harvested from the shoots of 4 week old plants, frozen in liquid N2 in 2 ml eppendorf tubes with metal beads, and ground in a tissue lyzer (MM400, Retsch). Ground plant material was resuspended in 500 μl extraction buffer (150 mM NaCl, 50 mM Tris-HCl pH 7.5, 10 % [v/v] glycerol, 1 % [v/v] Triton X-100, 5 mM DTT) containing a protease inhibitor cocktail (P9599, Sigma, 1 tablet / 20 ml) and centrifuged for 30 min and 1700 × g at 4°C. Protein concentrations were estimated in Bradford protein assays. Around 100 μg protein was mixed with 2x SDS loading buffer, boiled at 95 ^o^C for 10 min, and separated on 8 % SDS-PAGE gels. Anti-GFP antibody coupled with horse radish peroxidase (HRP, Miltenyi Biotec) at 1:2000 dilution was used to detect eGFP/mCit tagged protein constructs.

### Structural diagrams

for the crystals structures of *E. coli* PPK1 (http://rcsb.org PDB-ID 1XDO) (Zhu *et al.*, 2005) and PPX (PDB-ID 1U6Z) (Alvarado *et al.*, 2006) were prepared using Pymol (https://sourceforge.net/projects/pymol/).

## Acknowledgments

The Chlamydomonas WT 2C (mt^+^) strain was kindly provided by Michel Goldschmidt-Clermont, Department of Botany and Plant Biology, University of Geneva. We thank Michel Goldschmidt-Clermont and Chloe Laligne for help with Chlamydomonas cell culture and Yvon Jaillais, Daniel Couto and Michel Goldschmidt-Clermont for critical reading of the manuscript. This work was supported by an ERC starting grant from the European Research Council under the European Union’s Seventh Framework Programme (FP/2007-2013) / ERC Grant Agreement n. 310856 and the Howard Hughes Medical Institute International Research Scholar Award, both to M.H..

---

**Figure S1.**
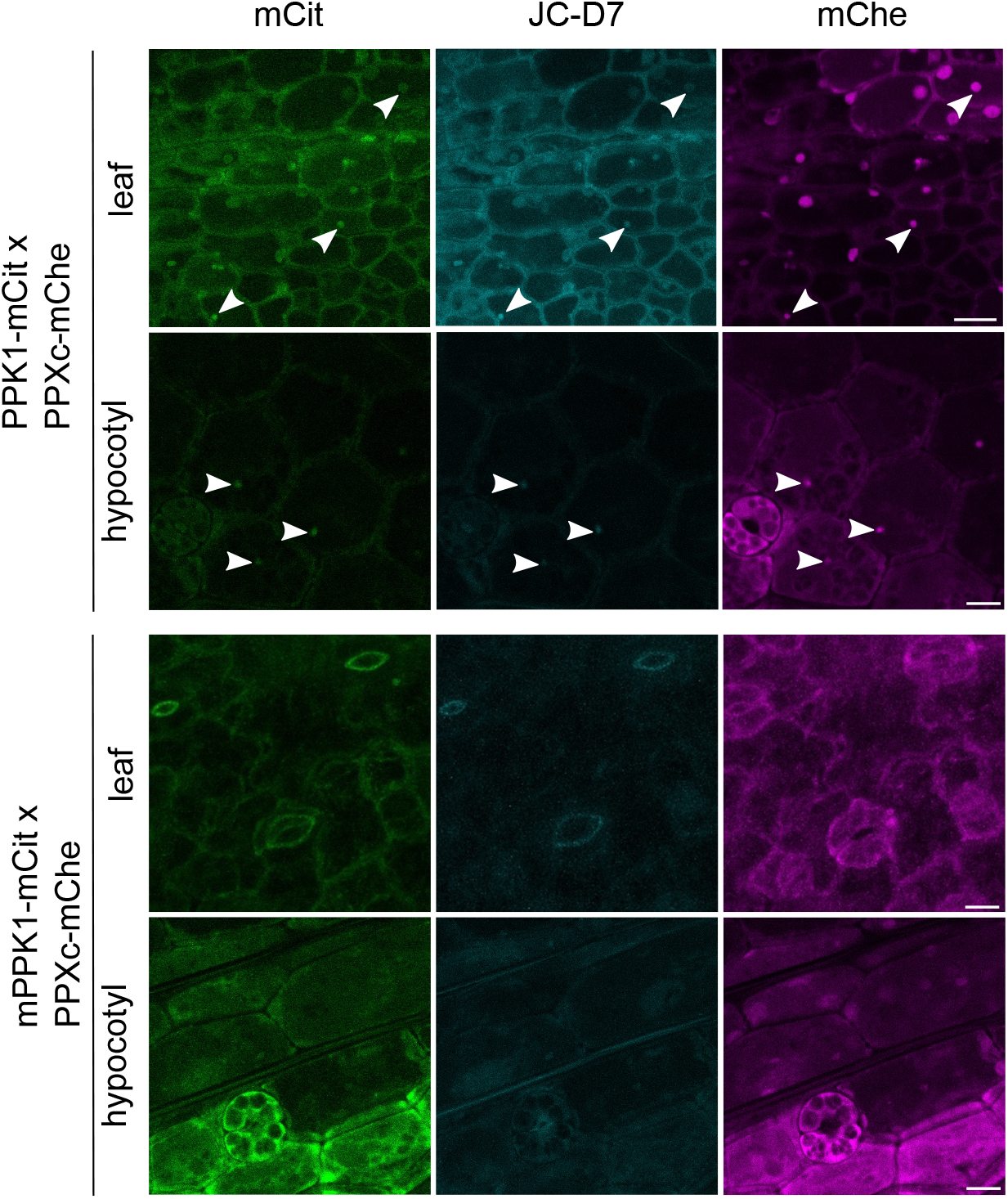
PolyP granules can be detected in fixed Arabidopsis cells expressing PPK1-mCit. Arabidopsis seeds from PPXc-mChe / PPK1-mCit (upper panel) or PPXc-mChe / mPPK1-mCit (lower panel) double transgenic lines were germinated on ^1/2^MS medium. 9 d old seedlings were fixed by para-formaldehyde and subsequently stained with JC-D7. Arrow heads highlight structures where PPXc-mChe (magenta) and active PPK1-mCit (green) co-localize with JC-D7 (cyan) stained polyP granules in leaf and hypocotyl cells. No co-localization or JC-D7 staining was observed in mPPK1-mCit expressing control lines. Scale bars = 10 μm.

**Figure S2.**
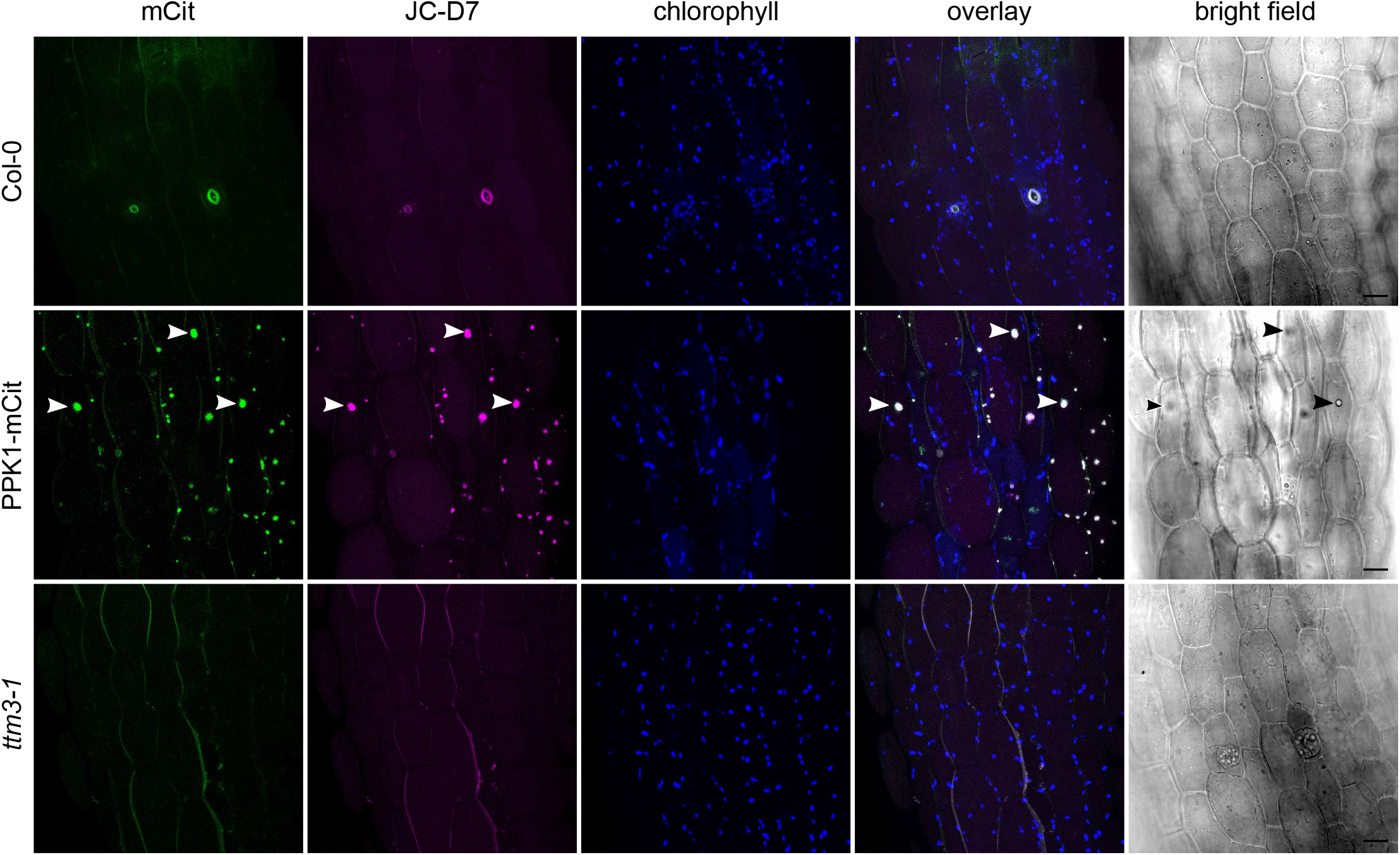
JC-D7 fails to stain specific structures in hypocotyl cells of Col-0 and *ttm3-1* plants. 9 d old Col-0 and *ttm1-3* seedlings grown on ^1/2^MS plates were stained by JC-D7 and imaged by confocal scanning microscopy, the PPK1-mCit line is shown as a control alongside. White arrow heads show PPK1-mCit (green) and JC-D7 (magenta) stained polyP granules in PPK1-mCit expressed hypocotyl cells, no specific JC-D7 staining was observed in Col-0 or *ttm3-1* hypocotyl cells. Scale bars = 20 μm.

**Figure S3.**
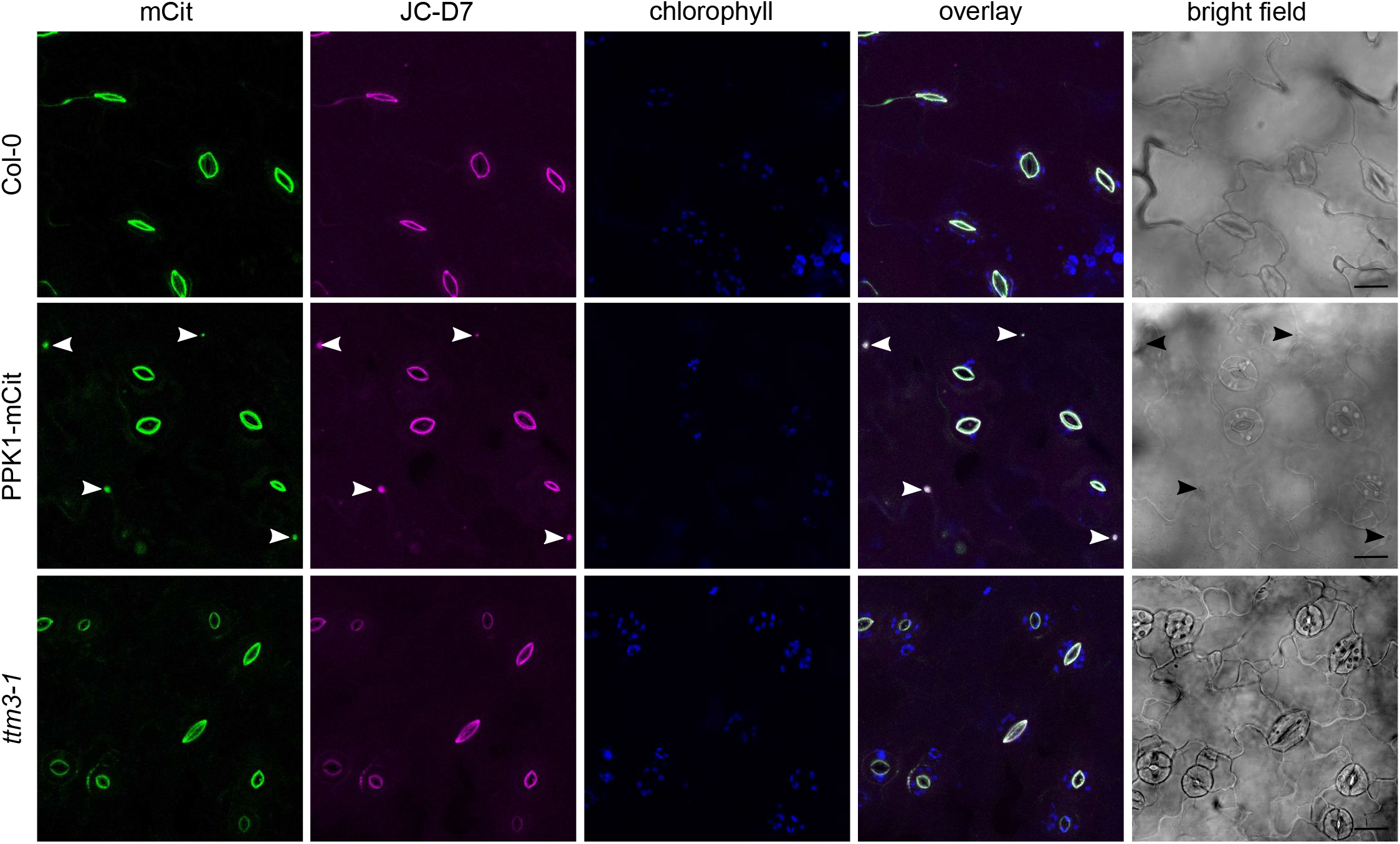
JC-D7 fails to stain specific structures in fixed hypocotyl cells of Col-0 and *ttm3-1* plants. Samples were grown and imaged as described in Figure S2, using a fixation protocol specified in Experimental Procedures. Scale bars = 20 μm.

**Figure S4.**
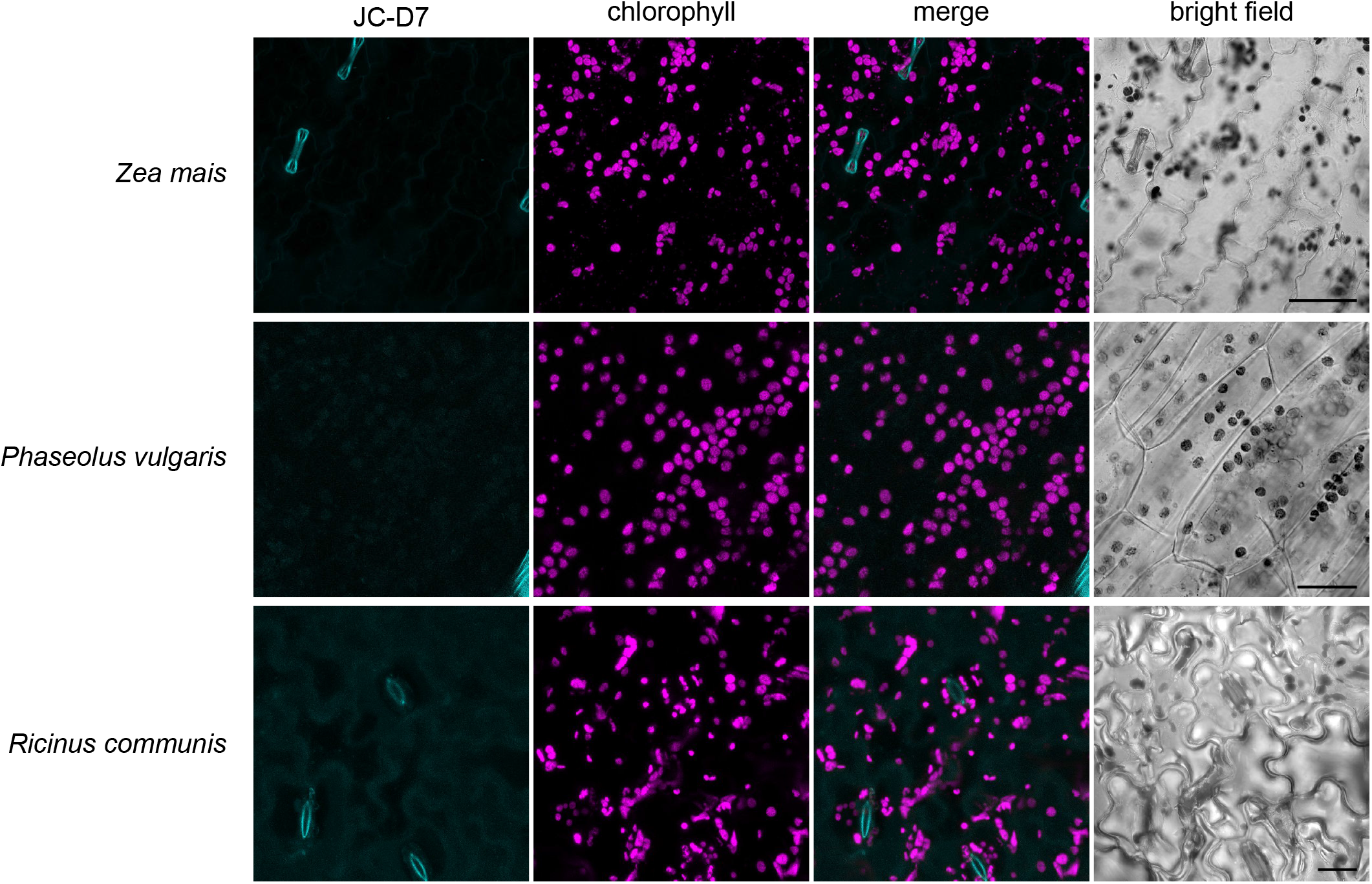
No JC-D7 stained polyP granules could be detected in epidermal cells of mono- and dicotyledonous plants. Epidermal cells from *Zea mais, Phaseolus vulgaris* and *Ricinus communis* were stained using the JC-D7 dye. No specific polyP signal was detected by confocal scanning microscopy.

## Notes

#### Summary of Updates

Species name corrected, figure legend corrected.

